# The role of epistasis in amikacin, kanamycin, bedaquiline, and clofazimine resistance in *Mycobacterium tuberculosis* complex

**DOI:** 10.1101/2021.05.07.443178

**Authors:** Roger Vargas, Luca Freschi, Andrea Spitaleri, Sabira Tahseen, Ivan Barilar, Stefan Niemann, Paolo Miotto, Daniella Maria Cirillo, Claudio U. Köser, Maha R. Farhat

**Affiliations:** Department of Systems Biology, Harvard Medical School, Boston, USA; Department of Biomedical Informatics, Harvard Medical School, Boston, USA; Emerging Bacterial Pathogens Unit, IRCCS San Raffaele Scientific Institute, Milan, Italy; National TB Reference laboratory, National TB Control Program, Islamabad, Pakistan; German Center for Infection Research, Partner site Hamburg-Lübeck-Borstel-Riems, Borstel, Germany; Molecular and Experimental Mycobacteriology, Research Center Borstel, Borstel, Germany; Department of Genetics, University of Cambridge, Cambridge, UK; Pulmonary and Critical Care Medicine, Massachusetts General Hospital, Boston, USA

## Abstract

Antibiotic resistance among bacterial pathogens poses a major global health threat. *M. tuberculosis* complex (MTBC) is estimated to have the highest resistance rates of any pathogen globally. Given the slow growth rate and the need for a biosafety level 3 laboratory, the only realistic avenue to scale up drug-susceptibility testing (DST) for this pathogen is to rely on genotypic techniques. This raises the fundamental question of whether a mutation is a reliable surrogate for phenotypic resistance or whether the presence of a second mutation can completely counteract its effect, resulting in major diagnostic errors (i.e. systematic false resistance results). To date, such epistatic interactions have only been reported for streptomycin that is now rarely used. By analyzing more than 31,000 MTBC genomes, we demonstrated that *eis* C-14T promoter mutation, which is interrogated by several genotypic DST assays endorsed by the World Health Organization, cannot confer resistance to amikacin and kanamycin if it coincides with loss-of-function (LoF) mutations in the coding region of *eis*. To our knowledge, this represents the first definitive example of antibiotic reversion in MTBC. Moreover, we raise the possibility that *mmpR* (*Rv0678*) mutations are not valid markers of resistance to bedaquiline and clofazimine if these coincide with LoF mutation in the efflux pump encoded by *mmpS5* (*Rv0677c*) and *mmpL5* (*Rv0676c*).

## INTRODUCTION

Tuberculosis (TB) and its causative pathogen *Mycobacterium tuberculosis* complex (MTBC) is a major public health threat causing an estimated 10 million new cases of disease per year (World Health Organization, 2020). Antibiotic resistance in particular poses a problem to controlling the TB epidemic (World Health Organization, 2020). Owing to the inherently slow growth rate of MTBC, genotypic drug-susceptibility testing (DST) represents the only realistic option to inform the initial selection of the most appropriate treatment regimen (Mohamed et al., 2021). This raises the fundamental question of whether the effect and clinical interpretation of a marker for resistance depends on the presence of another mutation (i.e. epistasis) or whether the effect is universal.

Although it is known that the level of resistance conferred by resistance mutations in some genes can differ, the only well-understood epistatic mechanism that completely counteracts the effect of another mutation involves the *whiB7* (*Rv3197A*) regulatory gene (Ajileye et al., 2017; Castro et al., 2020; Gagneux, 2018). Specifically, the over-expression of the *whiB7* cannot confer streptomycin resistance in the vast majority of lineage 2 isolates because these have a loss-of-function (LoF) mutation in the *tap* (*Rv1258c*) efflux pump (Köser et al., 2013; Merker et al., 2020). Yet, because the use of streptomycin has been downgraded in the most recent treatment guidelines by the World Health Organization (WHO), the clinical relevance of this example is limited (Viney et al., 2021).

Using whole-genome sequencing (WGS) data for 31,440 isolates, we set out to survey systematically whether other markers of resistance to more important antibiotics may be affected by epistasis if they involve the over-expression of a non-essential drug resistance gene. First, we analyzed the alkyl-hydroperoxidase *ahpC* (*Rv2428*), the function of which is not fully elucidated but may act as a compensatory mechanism for isoniazid (INH) resistance caused by *katG* mutations (i.e. the soon-to-be WHO-endorsed Cepheid Xpert MTB/XDR assay interrogates *ahpC* promoter/upstream mutations) (World Health Organization, In press). Second, LoF mutations in the transcriptional repressor *mmpR* (*Rv0678*), which is sequenced by several commercial targeted next-generation sequencing (tNGS) assays being evaluated by WHO, confer cross-resistance to bedaquiline (BDQ) and clofazimine (CFZ) via the over-expression of the non-essential efflux pump encoded by *mmpS5* (*Rv0677c*) and *mmpL5* (*Rv0676c*) (Kadura et al., 2020; Mohamed et al., 2021; Viljoen et al., 2017; Yamamoto et al., 2021). Third, four promoter/upstream mutations for the *eis* (*Rv2416c*) acetyltransferase are responsible for kanamycin (KAN) resistance, of which the C-14T mutation is due to be recognized by WHO as conferring cross-resistance to amikacin (AMK), the only aminoglycoside (AG) now recommended for the treatment of TB (World Health Organization, 2018, In press). In fact, the Xpert MTB/XDR already interprets this *eis* mutation accordingly, whereas the WHO-endorsed Hain GenoType MTBDR*sl* VER 2.0 (SL-LPA) assay will have to be updated accordingly. Finally, we included *whiB7* as it also regulates *eis* and, therefore, could theoretically confer cross-resistance to both AGs rather than just to KAN (World Health Organization, 2018, In press).

## RESULTS

### INH: *ahpC* upstream mutations in combination with *ahpC* LoF mutations

We observed 57 unique single nucleotide polymorphisms (SNPs) in the upstream region of *ahpC* (**Fig 1A**), of which 18 were homoplasic and occurred in at least five isolates, consistent with parallel evolution and known selection on this gene (**Table1, Supplementary Table 1**). We screened for frameshift indels, nonsense mutations, and mutations that abolish the start codon of *ahpC* given that these are the most likely types of mutations to confer a LoF phenotype. This yielded seven unique variants in eight isolates, of which just *ahpC* 323delC co-occurred with an upstream mutation in a single isolate (**Table 2, Supplementary Table 1**). This particular upstream mutation (i.e. G-88A **Table 1, Supplementary Table 1**) is a marker for MTBC lineage 3 and correlates with only a 3-fold increase in the expression of *ahpC*, potentially by creating a new Pribnow box (Chiner-Oms et al., 2019; Merker et al., 2020). As a result, this SNP is not considered to be a marker of resistance (i.e. the Xpert MTB/XDR was designed not to detect it, unlike adjacent mutations), which means that this is not an example of epistasis (World Health Organization, In press). Indeed, this double mutant was phenotypically susceptible to INH at the critical concentration (CC) of 0.1 mg/L in MGIT 960.

**Fig. 1.**
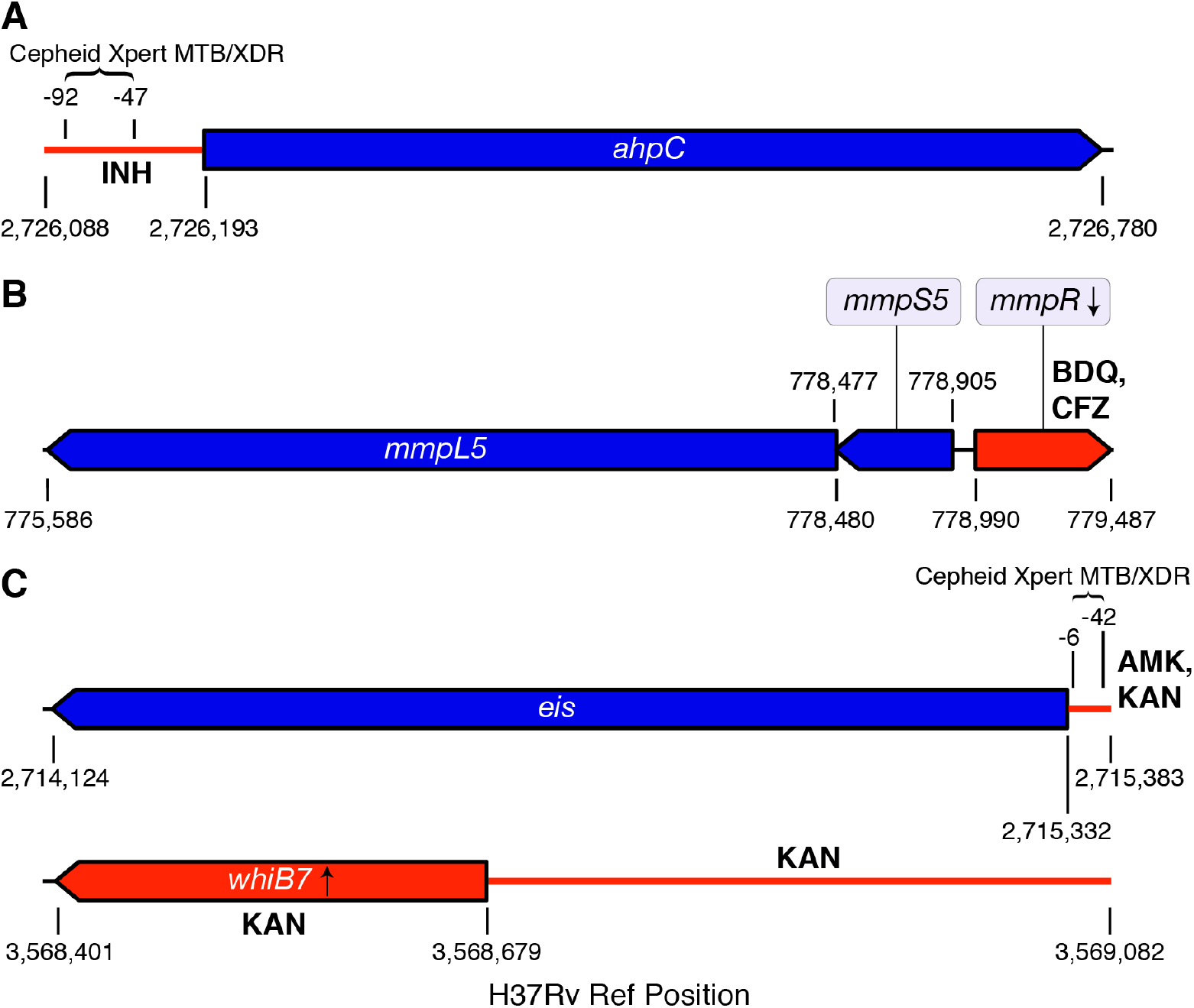
Genomic regions interrogated. The non-essential genes conferring resistance or compensating for resistance are shown in blue, whereas the corresponding regulatory regions or non-essential regulators are shown in red along with the relevant antibiotic(s). For each of the four regions, we screened for any type of mutation in the upstream regions and likely LoF mutations in the coding regions (i.e. frameshift indels, nonsense mutations, and mutations that abolish the start codon, including synonymous mutations at the start codons of *eis, mmpR*, and *whiB7* as these genes start with a valine). Unlike Cepheid, Hain has not disclosed the precise *eis* promoter region interrogated by its WHO-endorsed SL-LPA, which is why this information could not be included (Hain Lifescience, 2017).

**Table 1.** Mutations detected in a global sample of MTBC clinical isolate. Mutations that occur in our sample of 31,440 clinical isolates within the *mmpL5, mmpS5, mmpR, ahpC, eis, whiB7* coding sequences and *oxyR*-*ahpC, eis*-Rv2417c, *whiB7*-*uvrD2* intergenic regions (**Figure 1**). Mutations in **regulator** regions (*mmpR, oxyR*-*ahpC, eis*-Rv2417c, and *whiB7*-*uvrD2*) reported in this table were among the four most commonly detected variants in each region. Mutations in **regulated** regions (*mmpL5, mmpS5, ahpC, eis*, and *whiB7*) reported in this table were present in at least two MTBC sub-lineages. The full set of mutations detected within these genomic regions is reported in **Supplementary Table 1** (S=Synonymous, N=Non-Synonymous, I=Intergenic).

| Position | Variant Name | Gene Position | Mutation Type | Codon Position | # Isolates | # Sublineages |
| --- | --- | --- | --- | --- | --- | --- |
| 776210 | *mmpL5* C2271A | 2271 | N | Y757* | 2 | 2 |
| 777499 | *mmpL5* C982T | 982 | N | R328* | 2 | 2 |
| 777581 | *mmpL5* C900A | 900 | N | Y300* | 293 | 2 |
| 778086 | *mmpL5* 395insC | 395 | ins | 132 | 6 | 2 |
| 779127 | *mmpR* 138insG | 138 | ins | 46 | 5 | 4 |
| 779181 | *mmpR* 192-198delG | 192 | del | 64 | 20 | 4 |
| 779181 | *mmpR* 192-198insG | 192 | ins | 64 | 86 | 2 |
| 779407 | *mmpR* 418insG | 418 | ins | 140 | 6 | 1 |
| 2714753 | *eis* C580T | 580 | N | Q194* | 10 | 2 |
| 2715287 | *eis* 46insC | 46 | ins | 16 | 3 | 2 |
| 2715305 | *eis* G28T | 28 | N | E10* | 6 | 2 |
| 2715330 | *eis* G3A | 3 | S | V1V | 6 | 2 |
| 2715342 | *eis* G-10A | -10 *eis* | I |  | 293 | 18 |
| 2715344 | *eis* C-12T | -12 *eis* | I |  | 332 | 17 |
| 2715346 | *eis* C-14T | -14 *eis* | I |  | 181 | 19 |
| 2715369 | *eis* G-37T | -37 *eis* | I |  | 285 | 8 |
| 2726105 | *ahpC* G-88A | -88 *ahpC* | I |  | 3350 | 12 |
| 2726141 | *ahpC* C-52A | -52 *ahpC* | I |  | 91 | 10 |
| 2726141 | *ahpC* C-52T | -52 *ahpC* | I |  | 92 | 25 |
| 2726145 | *ahpC* G-48A | -48 *ahpC* | I |  | 85 | 20 |
| 3568487 | *whiB7* 193insG | 193 | ins | 65 | 3 | 3 |
| 3568488 | *whiB7* 192delC | 192 | del | 64 | 573 | 3 |
| 3568547 | *whiB7* 133delCA | 133 | del | 45 | 2 | 2 |
| 3568626 | *whiB7* 54delA | 54 | del | 18 | 61 | 2 |
| 3568779 | *whiB7* T-100C | -100 *whiB7* | I |  | 256 | 2 |
| 3568857 | *whiB7* C-178T | -178 *whiB7* | I |  | 73 | 2 |
| 3568921 | *whiB7* G-242C | -242 *whiB7* | I |  | 117 | 2 |
| 3569029 | *whiB7* A-350G | -350 *whiB7* | I |  | 249 | 3 |

**Table 2.** Co-occurrence of regulator resistance mutations and regulon LoF mutations. A list of antibiotic resistance mutations in **regulator** regions (*mmpR, oxyR*-*ahpC, eis*-Rv2417c, *whiB7*- *uvrD2*) that co-occur with LoF mutations in corresponding **regulated** regions (*mmpL5, mmpS5, mmpR, ahpC, eis, whiB7*) within our sample of 31,440 clinical isolates. A more detailed table can be found in **Supplementary Table 3**.

| A: Mutation in Regulator |  |  |  | B: Mutation in Regulated Gene |  |  |  | isolate info with co-occurring mutations |  |
| --- | --- | --- | --- | --- | --- | --- | --- | --- | --- |
| A type | A variant-name | A codon-pos | A num-isolates | B type | B variant-name | B codon-pos | B num-isolates | # isolates both mutations | isolate sublineages |
| INDEL | *mmpR* 192-198delG | 64 | 20 | INDEL | *mmpL5* 2028insA | 676 | 1 | 1 | 2.2.1.1.1(1) |
| INDEL | *mmpR* 192-198insG | 64 | 86 | INDEL | *mmpL5* 606delC | 202 | 83 | 82 | 4.11(82) |
| INDEL | *mmpR* 207insA | 69 | 1 | INDEL | *mmpL5* 1160insCGATG | 387 | 1 | 1 | 2.2.1(1) |
| SNP | *eis* C-14T |  | 181 | INDEL | *eis* 627insC | 209 | 1 | 1 | 2.2.1.1.1.i3(1) |
| SNP | *eis* C-14T |  | 181 | INDEL | *eis* 486insCT | 162 | 2 | 2 | 4.1.i1.1.1.1(2) |
| SNP | *eis* C-14T |  | 181 | INDEL | *eis* 473insT | 158 | 1 | 1 | 2.2.1.1.1.i3(1) |
| SNP | *eis* C-14T |  | 181 | INDEL | *eis* 448delA | 150 | 7 | 7 | 2.2.1.1.1.i3(7) |
| SNP | *eis* C-14T |  | 181 | INDEL | *eis* 400insG | 134 | 1 | 1 | 2.2.1.1.1(1) |
| SNP | *eis* C-14T |  | 181 | INDEL | *eis* 279delCGGCGATGCGT | 93 | 1 | 1 | 2.2.1.1.1.i3(1) |
| SNP | *eis* C-14T |  | 181 | SNP | *eis* G39A | W13* | 1 | 1 | 2.2.1.1.1(1) |
| SNP | *eis* C-14T |  | 181 | SNP | *eis* G38A | W13* | 1 | 1 | 2.2.1.1.1(1) |
| SNP | *eis* C-14T |  | 181 | INDEL | *eis* 16insC | 6 | 1 | 1 | 2.2.1.1.1(1) |
| SNP | *eis* C-14T |  | 181 | INDEL | *eis* 15insC | 5 | 1 | 1 | 2.2.1.1.1(1) |
| SNP | *eis* C-14T |  | 181 | SNP | *eis* G3A | V1V | 6 | 6 | 1.1.1.1.1(1) & 2.2.1.1.1.i3(5) |
| SNP | *ahpC* G-88A |  | 3350 | INDEL | *ahpC* 323delC | 108 | 1 | 1 | 3.1.1(1) |
| SNP | *whiB7* T-147C |  | 1 | INDEL | *whiB7* 192delC | 64 | 573 | 1 | 1.2.1.1.1(1) |
| INDEL | *whiB7* -214delG |  | 9 | INDEL | *whiB7* 192delC | 64 | 573 | 1 | 1.2.1.1.1(1) |
| INDEL | *whiB7* -316insC |  | 2 | INDEL | *whiB7* 192delC | 64 | 573 | 1 | 1.2.1.1.1(1) |

### BDQ/CFZ: *mmpR* LoF mutations in combination with *mmpS5*-*mmpL5* LoF mutations

We detected 91 fixed LoF variants in *mmpL5*-*mmpS5*-*mmpR*, of which 35 occurred in at least two isolates (**Fig 1B, Supplementary Table 1**). Frameshifts were most common (39/68) in *mmpL5*, followed by *mmpR* (21/68), and *mmpS5* (8/68). Each gene harbored frameshifts in isolates from at least three MTBC major lineages, indicating parallel evolution (**Supplementary Table 1**). The nonsense SNP *mmpL5* Y300* was observed in 293 isolates and in two genetically distinct lineages, and the insC at nt395 of *mmpL5* also occurred two in distant lineages (**Table 1, Supplementary Table 1**). The *mmpR* delG in the homopolymer (HP) nt192-198 was observed in 20 isolates from three major lineages, whereas insG in the same HP was observed in 86 isolates from two major lineages (**Table 1, Supplementary Table 1**). Noting the frequency of frameshifts in the homopolymer region of *mmpR*, we investigated non-fixed frameshift variants (*i*.*e*. that had within-sample allele frequencies of 10-75%) and recorded which isolates had >100x coverage of *mmpR, mmpS5*, and *mmpL5*. Frameshift variants at low to intermediate allele frequency were rare and occurred in a total of six isolates (3/7435 in *mmpR*, 1/8949 in *mmpS5*, and 2/6217 in *mmpL5*). Two of these isolates had the frameshift insG in the aforementioned *mmpR* HP at 66% and 71% allele frequencies (**Fig. 2, Supplementary Table 2**).

**Fig. 2.**
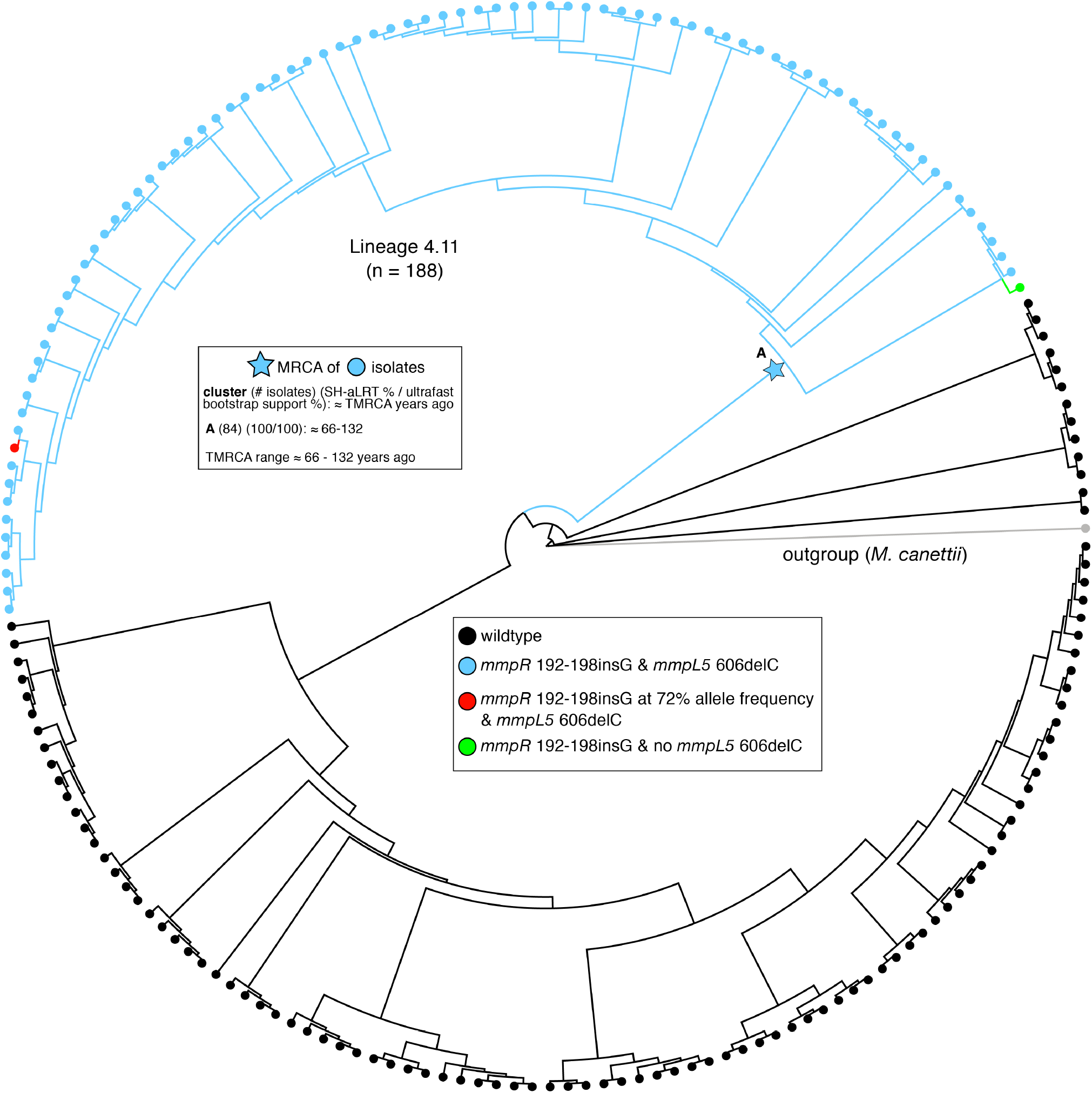
Phylogeny of 188 sub-lineage 4.11 isolates. Isolates with both the *mmpR* 192-198insG and *mmpL5* 606delC variants are colored in blue (n=82), isolates carrying neither variant are colored in black (n=104). One isolate had the *mmpR* 192-198insG frameshift but not the *mmpL5* 606delC frameshift (green). Another isolate had the *mmpL5* 606delC frameshift and 72% of reads supporting the *mmpR* 192-198insG frameshift (red). Time to most recent common ancestor (TMRCA) estimates for the group of isolates with the *mmpR* 192-198insG and *mmpL5* 606delC variants is given in the upper left.

Three different LoF mutations in *mmpR* coincided with a LoF mutation in *mmpL5* (**Fig. 2**). Of those, insG in the HP nt192-198, which had been repeatedly demonstrated to confer BDQ and CFZ resistance during *in vitro* selection experiments and patient treatment, occurred in 82 isolates, whereas the other two double mutations were observed in only a single isolate, respectively (Andres et al., 2020; de Vos et al., 2019; Ghodousi et al., 2019; Peretokina et al., 2020; Sonnenkalb et al., 2021; Zhang et al., 2015). All of the former 82 double-LoF mutants belonged to a monophyletic group within sub-lineage 4.11 that was mostly multi-drug resistant (53 of 59 with known phenotypic data). Most double-LoF mutants were isolated in Lima, Peru, between 1997 and 2012 and represented 43% (82/188) of the isolates from the sub-lineage 4.11 in our dataset (**Fig. 2, Table 2, Supplementary Table 3-4**). Among the 84 isolates with co-occurance of *mmpR* and *mmpL5* LoF, there were no SNPs in the other BDQ resistance locus, *atpE*.

We constructed a phylogeny of all 188 MTBC sub-lineage 4.11 isolates and to study how the LoF mutations in *mmpR* and *mmpS5-mmpL5* evolved (**Fig. 2**). The majority of isolates with the *mmpR* or *mmpL5* frameshifts harbored both (82/84) but, based on the topology of the tree, we were unable determine which of the two frameshifts arose first. Consequently, we could only date the common most recent common ancestor (MRCA). We approximated the age of the MRCA at 66-132 years prior to sampling *i*.*e*. well before the use of BDQ or CFZ in treatment regimens for TB in the pre-antibiotic era and likely before the introduction of thioacetazone, which is also exported by *mmpS5-mmpL5* (Halloum et al., 2017; Ma et al., 2010).

### KAN: *eis* upstream mutations in combination with *eis* LoF mutations

We observed 23 unique LoF mutations upstream of *eis* (**Fig. 1C**), of which ten were homoplasic and occurred in at least five isolates (**Supplementary Table 1**). As expected, the classical G-37T, C-14T, C-12T, and G-10A mutations, which are known to confer KAN resistance based on allelic exchange and/or complementation experiments, were most frequent (**Table 1**) (Pholwat et al., 2016; World Health Organization, 2018; Zaunbrecher et al., 2009). Specifically, 881 isolates with either *eis* G-37T, C-12T, or G-10A and 179 isolates with *eis* C-14T did not have any of the other key AG resistance mutations in *rrs* (i.e. A1401G, C1402T, or G1484T, see **Supplementary Table 5**) (World Health Organization, In press).

We identified 30 unique LoF mutations in *eis*, of which five were homoplasic and occurred in at least five isolates (**Table 1, Supplementary Table 1**). These LoF never coincided with *eis* G-37T, C-12T, or G-10A, whereas this was the case for 21 *eis* C-14T mutants (i.e. 13 isolates with indels, six with a G3A synonymous change that abolished the valine start codon, and two with nonsense mutations) (**Table 2, Supplementary Table 3**). MIC data were available for five of these *eis* double mutants, which confirmed that they were susceptible to KAN whereas seven *eis* C-14T control isolates with a wild-type *eis* coding region were KAN resistant (**Fig. 3, Table 3**). The corresponding AMK MIC data mirrored the results for KAN.

**Fig. 3.**
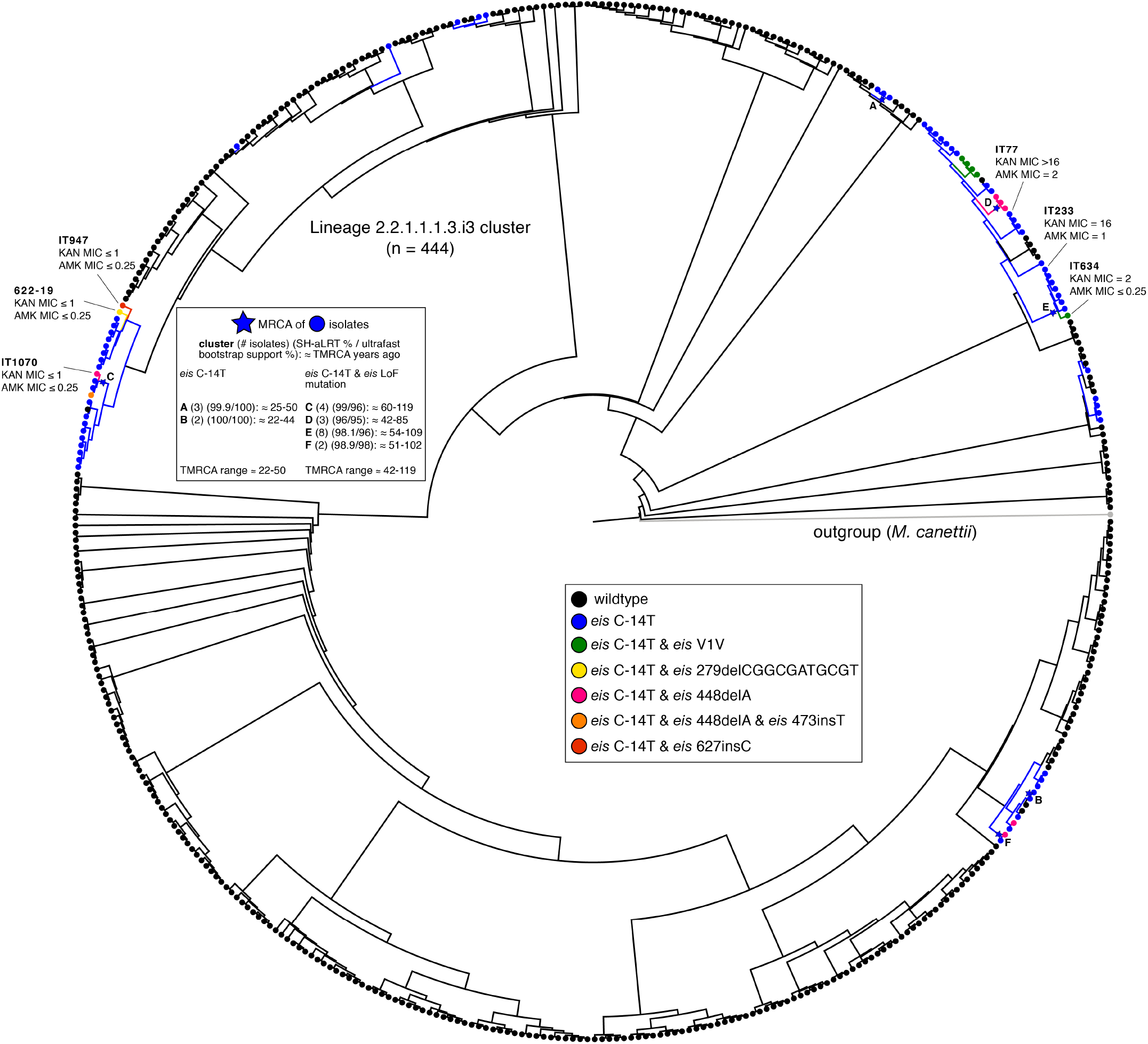
Phylogeny of 444 sub-lineage 2.2.1.1.1.3.i3 isolates. Isolates with the *eis* C-14T promoter SNP and no LoF variants in *eis* are colored in blue (n=61), whereas isolates carrying both the *eis* C-14T promoter SNP and a LoF mutation in *eis* are colored according to the legend (n=14). TMRCA estimates for groups of isolates with the *eis* C-14T promoter SNP are given in the upper left. The MICs (mg/L) for isolates from this sub-lineage are also included (MICs for isolates from other sub-lineages can be found in **Table 3**).

**Table 3.**
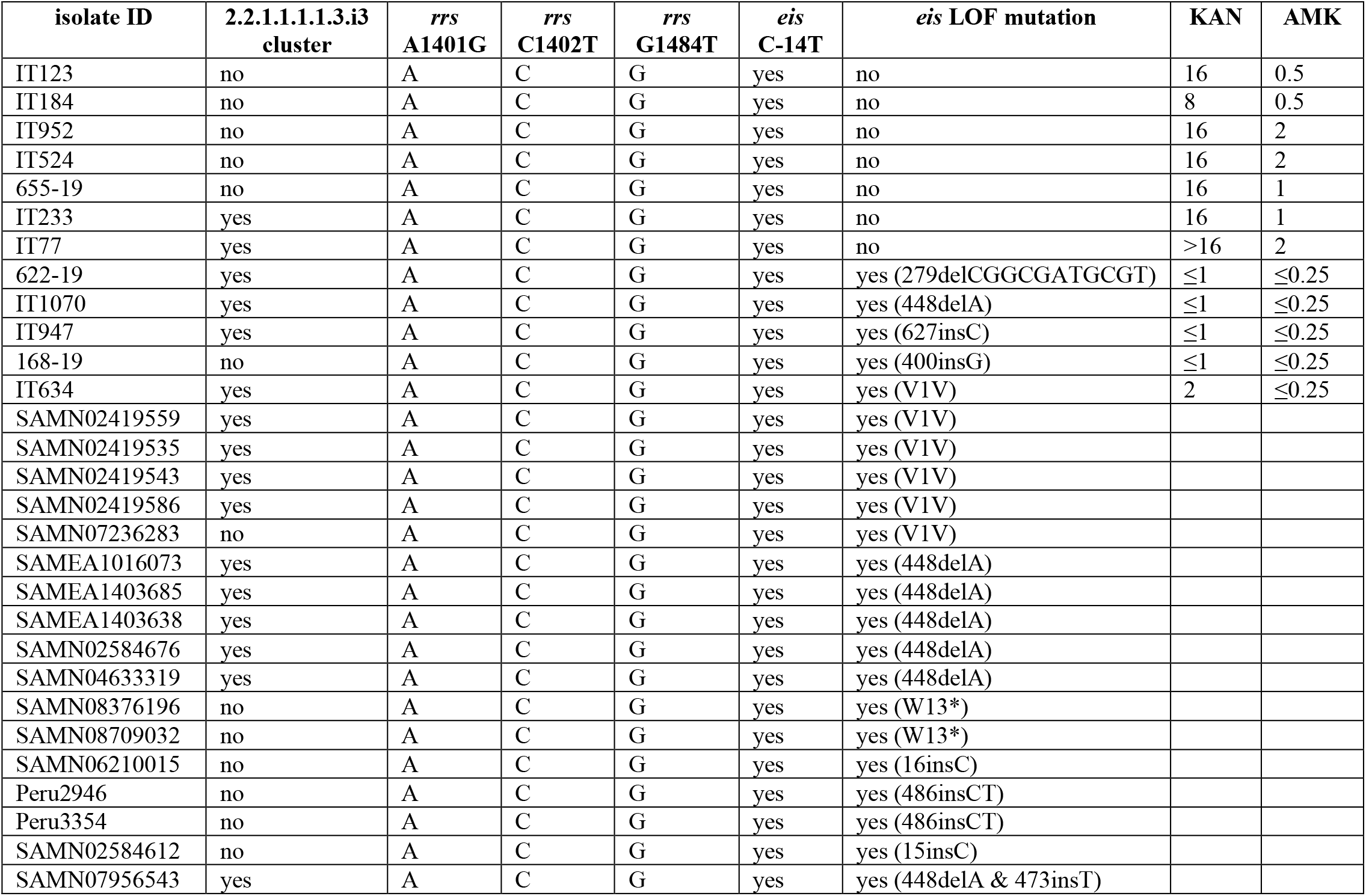
Overview of *eis* C-14T isolates with *eis* LoF mutations and corresponding MICs for KAN and AMK. All 29 double mutants lacked the three classical AG resistance mutations in *rrs*. Isolates that are part of the 2.2.1.1.1.1.3.i3 cluster are shown in Fig 3. MICs (mg/L) were measured using either the UKMYC5 or UKMYC6 broth microdilution plates by Thermo Fisher (Fowler and CRyPTIC Consortium, 2021). The provisional CCs for KAN and AMK for these plates are 4 and 1 mg/L, respectively. Unlike for KAN, *eis* C-14T only has a modest effect on the MIC of AMK (i.e. the MIC distribution of this mutation spans the CC when the efflux pump is active, which is in line with data from other media) (Gygli et al., 2019; Pholwat et al., 2016; World Health Organization, 2018; Zaunbrecher et al., 2009). Consequently, KAN is a more sensitive agent to detect an inactive efflux pump than AMK. More details can be found in **Supplementary Table 6**.

The most common MTBC sub-lineage with the *eis* double mutants was 2.2.1.1.1.i3 (14/22 isolates). Of the 31,440 isolates, 444 were belonged to this sub-lineage and clustered closely based on their pairwise SNP distance. The phylogeny of these 444 isolates showed that *eis* C-14T arose more than nine times independently (**Fig. 3**). We approximated the MRCA for the six group of isolates that had high bootstrap support. For the two groups of isolates that only harbored the *eis* C-14T mutation the MRCA was dated to 22-50 years ago, in line with KAN introduction into clinical use in approximately 1958 (Ektefaie et al., 2021). Of the 75 isolates with the *eis* C-14T promoter mutation, 14 also harbored an *eis* LoF mutation that arose nine times independently (**Fig. 3, Table 2**). In each instance, the LoF variant emerged within a clade of *eis* C-14T mutants suggesting that it appeared later in time. We compared the MRCA of the clades with double mutants to those with *eis* C-14T only. We found the MRCA of the double mutants to be older on average, suggesting that time and possibly fluctuating evolutionary pressures are needed for LoF to develop in an *eis* C-14T background.

### KAN: *whiB7* upstream mutations in combination with *whiB7* or *eis* LoF mutations

We found 116 unique SNPs upstream of *whiB7* (**Fig. 1C**), of which 8 were homoplasic and occurred in at least five isolates (**Table 1, Supplementary Table 1**). We identified 10 unique LoF mutations in *whiB7*, two of which (nt193insG & nt133delCA) evolved repeatedly, across 657 isolates. The most frequent mutation (nt192delC) occurred in 573 sub-lineage 1.2.1.1 isolates, which was in agreement with earlier findings (Merker et al., 2020). This was the only LoF mutation in *whiB7* to coincide with an upstream mutation (i.e. in three isolates in total, each with a different upstream mutation) (**Table 2, Supplementary Table 3**). Because none of these upstream mutations had been described in the literature, it was unclear whether these represented potential examples of epistasis (Heyckendorf et al., 2018; Reeves et al., 2013). Finally, no LoF mutations in *eis* were found in isolates harboring mutations upstream of *whiB7*.

## DISCUSSION

Although our analysis did not yield any strong evidence for epistasis involving *ahpC* or *whiB7*, our finding that epistasis is possible due to LoF mutations in *eis* is not only relevant for AGs but has wider implications for the interpretation of sequencing data. First, with the exception of two synonymous mutations in *aftA* (*Rv3792*) and *fabG1* (*Rv1483*) that confer ethambutol and ethionamide/INH resistance respectively by creating alternative promoters, synonymous mutations are typically excluded *a priori* from the analysis of WGS data (Ando et al., 2014; Safi et al., 2013). We demonstrated that this assumption is not sound for start codons given that only one of the four triplets encoding valine can act as a start codon (i.e. GTC). Second, evidence for epistasis argues strongly that multivariate prediction approaches are needed for accurate resistance prediction from sequencing data.

It is notable that *eis* LoF mutations coincided only with *eis* C-14T mutants, even though isolates with *eis* G-37T, C-12T, and G-10A without any AG resistance mutations in *rrs* were almost five times more frequent in our dataset (**Supplementary Table 5**). We hypothesize that because *eis* C-14T leads to a greater up-regulation of *eis* than the other three mutations, this comes at a fitness cost, unless selective antibiotic pressure is maintained (Pholwat et al., 2016; Sanz-García et al., 2019; World Health Organization, 2018; Zaunbrecher et al., 2009). Indeed, molecular dating and the topology of the in tree (**Fig. 3**) suggested that the LoF mutations arose independently on multiple occasions after the acquisition of *eis* C-14T. To our knowledge, this represents the strongest evidence to date for genotypic reversion from resistance to a susceptible phenotype for MTBC (Richardson et al., 2009). We would like to stress, however, that even for AMK this is a rare phenomenon, given that of the 179 isolates that harbored *eis* C-14T without any AG resistance mutations in *rrs* only 12% had concomitant *eis* LoF mutations (**Table 2, Supplementary Table 5**). In other words, the cautious approach would be to still interpret *eis* C-14T as a marker for AMK resistance to construct a relevant regimen, unless there is strong evidence that a particular isolate is affected by epistasis (e.g. unlike the SL-LPA, the tNGS assays by ABL and Deeplex actually interrogate part of the *eis* coding region) (Mohamed et al., 2021; World Health Organization, 2018). This initial treatment decision may then have to be adjusted based on the phenotypic DST result.

Because we did not have BDQ or CFZ MICs for any of the *mmpR*/*mmpL5* double-LoF mutants, it remains to be determined whether these are examples of epistasis (in the case of the Peruvian cluster, this would be unrelated to antibiotic pressure, unlike for *eis*). We note, however, that indirect evidence exists that is in line with this prediction. Villellas et al. reported the BDQ MICs and *mmpR* sequence results for baseline isolates from the C208 phase 2b trial of BDQ, which featured five isolates with the same *mmpR* frameshift that we observed in the Peruvian cluster (**Fig. 2**) (Diacon et al., 2014; Villellas et al., 2017). Three of the trial mutants were from South Africa and had 7H11 MICs of 0.25-1 mg/L (i.e. ≥CC of 0.25 mg/L and, thus, consistent with a functional efflux pump and resistant phenotype if an area of technical uncertainly is set at 0.25 mg/L, as previously proposed) (Beckert et al., 2020; Nimmo et al., 2020; Villellas et al., 2017). By contrast, even the lowest BDQ concentration tested (i.e. 0.008 mg/L) inhibited the growth of the remaining two trial mutants that were isolated in 2009 in Lima, Peru (N. Lounis, personal communication). Given that the Peruvian double-LoF cluster from this study was isolated in the same city during the same period, it is possible that the latter two trial isolates were from this cluster, although this remains to be confirmed using WGS data and, ideally, repeat MIC testing to exclude experimental error.

The possibility of epistasis underlines the need for comprehensive microbiological workup of the ongoing clinical trials of BDQ. *mmpR* as well as *mmpS5-mmpL5* and the corresponding inter-genic region have to be analyzed along with standardized MIC testing using an on-scale quality control strain for both BDQ and CFZ (Kaniga et al., 2016; Schön et al., 2020, 2019). We recommend that discordances between genotypic and phenotypic DST results are confirmed by retesting and, where warranted, followed up with specialized testing. For example, the two aforementioned Peruvian results may actually be hyper-susceptible to BDQ and CFZ (i.e. lower concentrations would have to be tested to determine the MIC endpoint) (Merker et al., 2020). If confirmed, this should also apply to *mmpS5*-*mmpL5* LoF mutants with wild-type *mmpR* (e.g. just over half of the lineage 1.1.1.1 isolates in our dataset had a nonsense mutation in *mmpL5*) and may have implications for the ongoing trials of TBAJ-876, TBAJ-587, TBI-166, and OPC-167832 as these agents are also exported by this pump (Hariguchi et al., 2020; Xu et al., 2021, 2019).

## MATERIALS AND METHODS

### Sequence Data

We initially downloaded raw Illumina sequence data for 33,873 clinical isolates from NCBI (Benson et al., 2008). We identified the BioSample for each isolate and downloaded all of the associated Illumina sequencing runs. Isolates had to meet the following quality control measures for inclusion in our study: (i) at least 90% of the reads had to be taxonomically classified as belonging to MTBC after running the trimmed FASTQ files through Kraken (Wood and Salzberg, 2014) and (ii) at least 95% of bases had to have coverage of at least 10x after mapping the processed reads to the H37Rv reference genome (Genbank accession: NC_000962).

### Illumina Sequencing FastQ Processing and Mapping to H37Rv

The raw sequence reads from all sequenced isolates were trimmed with version 0.20.4 Prinseq (settings: -min_qual_mean 20) (Schmieder and Edwards, 2011) and then aligned to H37Rv with version 0.7.15 of the BWA mem algorithm using the -M settings (Li and Durbin, 2009). The resulting SAM files were then sorted (settings: SORT_ORDER = coordinate), converted to BAM format, and processed for duplicate removal with version 2.8.0 of Picard (http://broadinstitute.github.io/picard/) (settings: REMOVE_DUPLICATES = true, ASSUME_SORT_ORDER = coordinate). The processed BAM files were then indexed with Samtools (Li et al., 2009). We used Pilon (settings: --variant) on the resulting BAM files to generate VCF files that contained calls for all reference positions corresponding to H37Rv from pileup (Walker et al., 2014).

### Empirical Score for Difficult-to-Call Regions

We assessed the congruence in variant calls between short-read Illumina data and long-read PacBio data for a set of isolates that underwent sequencing with both technologies. Using 31 isolates for which both Illumina and a complete PacBio assembly were available, we evaluated the empirical base-pair recall (EBR) of all base-pair positions of the H37rv reference genome. For each sample, the alignments of each high confidence genome assembly to the H37Rv genome were used to infer the true nucleotide identity of each base pair position. To calculate the empirical base-pair recall, we calculated what percentage of the time our Illumina based variant calling pipeline, across 31 samples, confidently called the true nucleotide identity at a given genomic position. If Pilon variant calls did not produce a confident base call (*Pass*) for the position, it did not count as a correct base call. This yields a metric ranging from 0.0–1.0 for the consistency by which each base-pair is both confidently and correctly sequenced by our Illumina WGS based variant calling pipeline for each position on the H37Rv reference genome. An H37Rv position with an EBR score of x% indicates that the base calls made from Illumina sequencing and mapping to H37Rv agreed with the base calls made from the PacBio *de novo* assemblies in x% of the Illumina-PacBio pairs. We masked difficult-to-call regions by dropping H37Rv positions with an EBR score below 0.9 as part of our variant calling procedure. Full details on the data and methodology can be found elsewhere (Vargas et al., 2021).

### Variant Calling

#### SNP Calling

To prune out low-quality base calls that may have arisen due to sequencing or mapping error, we dropped any base calls that did not meet any of the following criteria: (i) the call was flagged as *Pass* by Pilon, (ii) the mean base quality at the locus was >20, (iii) the mean mapping quality at the locus was >30, (iv) none of the reads aligning to the locus supported an insertion/deletion (indel), (v) a minimum coverage of 20 reads at the position, and (vi) at least 75% of the reads aligning to that position supported 1 allele (using the *INFO*.*QP* field which gives the proportion of reads supporting each base weighted by the base and mapping quality of the reads, *BQ* and *MQ* respectively, at the specific position). A base call that did not meet all filters (i) – (vi) was inferred to be low-quality/missing.

#### Indel Calling

To prune out low-quality indel variant calls, we dropped any indel that did not meet any of the following criteria: (i) the call was flagged as *Pass* by Pilon, (ii) the maximum length of the variant was 10bp, (iii) the mean mapping quality at the locus was >30, (iv) a minimum coverage of 20 reads at the position, and (v) at least 75% of the reads aligning to that position supported the indel allele (determined by calculating the proportion of total reads *TD* aligning to that position that supported the insertion or deletion, *IC* and *DC* respectively). A variant call that met filters (i), (iii), and (iv) but not (ii) or (v) was inferred as a high-quality call that did not support the indel allele. Any variant call that did not meet all filters (i), (iii), and (iv) was inferred as low-quality/missing.

#### Intermediate Allele Frequency Indel Calling

To call indel variants in which the indel allele was detected at an intermediate frequency, we made the following modification to the *indel Calling* filters outlined above. Filter (v) above is replaced with the following two filters: (v-i) at least 10% but less than 75% of the reads aligning to that position supported the indel allele and (v-ii) at least 10 reads support the indel allele. The *mmpR* analysis was restricted to isolates with 100x coverage across ≥99% of the gene.

### SNP Genotypes Matrix

We detected SNP sites at 899,035 H37Rv reference positions (of which 64,950 SNPs were not biallelic) among our global sample of 33,873 isolates. We constructed a 899,035×33,873 genotypes matrix (coded as 0:A, 1:C, 2:G, 3:T, 9:Missing) and filled in the matrix for the allele supported at each SNP site (row) for each isolate, according to the *SNP Calling* filters outlined above. If a base call at a specific reference position for an isolate did not meet the filter criteria that allele was coded as *Missing*. We excluded 20,360 SNP sites that had an EBR score <0.90, another 9,137 SNP sites located within mobile genetic element regions (e.g. transposases, intergrases, phages, or insertion sequences) (Comas et al., 2010; Vargas et al., 2021), then 31,215 SNP sites with missing calls in >10% of isolates, and 2,344 SNP sites located in overlapping genes (coding sequences). These filtering steps yielded a genotypes matrix with dimensions 835,979×33,873. Next, we excluded 1,663 isolates with missing calls in >10% of SNP sites yielding a genotypes matrix with dimensions 835,979×32,210. We used an expanded 96-SNP barcode to type the global lineage of each isolate in our sample (Freschi et al., 2020). We further excluded 325 isolates that either did not get assigned a global lineage, assigned to more than one global lineage, or were typed as lineage 7. We then excluded 41,760 SNP sites from the filtered genotypes matrix in which the minor allele count = 0 which resulted in a 794,219×31,885 matrix. To provide further MTBC lineage resolution on the lineage 4 isolates, we required an MTBC sub- lineage call for each lineage 4 isolate. We excluded 457 isolates typed as global lineage 4 but had no further sub-lineage calls and then again excluded 11,654 SNP sites from the filtered genotypes matrix in which the minor allele count=0. The genotypes matrix used for downstream analysis had dimensions 782,565×31,428, representing 782,565 SNP sites across 31,428 isolates. The global lineage (L) breakdown of the 31,428 isolates was: L1=2,815, L2=8,090, L3=3,398, L4=16,931, L5=98, L6=96.

### Indel Genotypes Matrix

We detected 53,167 unique indel variants within 50,576 H37Rv reference positions among our global sample of 33,873 isolates. We constructed a 53,167×33,873 genotypes matrix (coded as 1:high quality call for the indel allele, 0:high quality call not for the indel allele, 9:Missing) and filled in the matrix according to whether the indel allele was supported for each indel variant (row) for each isolate, according to the *Indel Calling* filters outlined above. If a variant call at the reference position for an indel variant did not meet the filter criteria that call was coded as *Missing*. We excluded 2,006 indel variants that had an EBR score <0.90, another 694 indel variants located within mobile genetic element regions, then 207 indel variants located in overlapping genes (coding sequences). These filtering steps yielded a genotypes matrix with dimensions 50,260×33,873. Next, we excluded any isolate that was dropped while constructing the SNP genotypes matrix to retain the same 31,428 isolates as described above. The genotypes matrix used for downstream analysis had dimensions 50,260×31,428.

### Mixed Allele Frequency Indel Genotypes Matrix

After following the same filtering steps outlined above under *Indel Genotypes Matrix*, we detected 7,731 unique indel variants in our filtered sample of 31,428 isolates in which at least one isolate supported each indel variant at an intermediate allele frequency (10%≤AR<75%). We constructed a 7,731×31,428 genotypes matrix (coded as 0:high quality call not for the indel allele, -9:Missing, or 10-74:the % of reads supporting the indel allele) and filled the matrix according to whether the indel allele was supported at an intermediate allele frequency for each indel variant (row) for each isolate, according to the *Intermediate Allele Frequency Indel Calling* filters outlined above. To determine the limit of detection for indels that might be present at lower allele frequencies, we calculated the number of isolates in our sample that have 100x coverage in ≥99% of the locus for *mmpR* (7,435), *mmpS5* (8,949), and *mmpL5* (6,217) (**Supplementary Table 2)**. We retained only frameshift indels yielding a genotypes matrix with dimensions 5,925×31,428 and interrogated only the *mmpR* - *mmpS5* - *mmpL5* chromosomal region for the presence of mixed indels (**Supplementary Table 2**).

### Inclusion and Processing of 12 *eis* C-14T mutants with AG MICs

We added 12 clinical *eis* C-14T mutants to the dataset, for which we had KAN and AMK MICs and some of which had a LoF mutation in *eis*. We processed the raw sequencing reads according to the methods described above to generate VCF files. We genotyped SNPs for these isolates at the 782,565 SNP sites and genotyped indels for the 50,260 indel variants previously identified using the same filters described above to construct 782,565×12 and 50,260×12 matrices, respectively.

During analysis, we observed that 3/12 isolates (IT947, 622-19 and 168-19) carried the *eis* C-14T promoter resistance mutation and no observed LoF mutation in *eis* but were phenotypically susceptible according to KAN MICs. Upon further inspection of the VCF files for these isolates, we found that all three isolates had a LoF mutation in *eis* that we originally did not detected per our variant calling methodology. We found that one isolate (622-19) had an 11bp deletion in *eis* which was not represented in the 50,260 indel variants since we restricted our analysis to indels ≤10bp and consequently was excluded from our 50,260×12 matrix. Each of the other two strains, IT947 and 168-19, had a different 1bp insertion in *eis* that was not identified in our original pool of 31,428 isolates, so it also was also not represented in the 50,260×12 matrix. We updated our variant call data by incorporating these newly identified variants (**Table 2, Supplementary 1, Supplementary Table 3**).

### Targeted Chromosomal Regions

We queried our SNP and indel matrices for the following types of mutations in the following regions of the H37Rv Reference Genome: [1] *<u>mmpR</u>*<u>-</u> *<u>mmpS5</u>*- *<u>mmpL5</u>*: the coding sequences for *mmpR* (778990 - 779487), *mmpS5* (778477 - 778905), and *mmpL5* (775586 - 778480) for nonsense SNVs (single nucleotide variant), frameshift indels, missense SNVs that abolish the start codon, and synonymous SNVs that abolish the start codon for *mmpR* which starts with a valine (we did not check for synonymous SNVs at the first codon for *mmpS5* or *mmpL5* because these coding sequences start with a methionine). [2] <u>upstream</u> *<u>ahpC</u>*<u>-</u> *<u>ahpC</u>*: the intergenic region *oxyR*- *ahpC* (2726088 - 2726192) for SNVs and indels, and the coding sequence for *ahpC* (2726193 - 2726780) for nonsense SNVs, frameshift indels, and missense SNVs that abolish the start codon. We did not check for synonymous SNVs at the first codon for *ahpC* because the coding sequence starts with a methionine (and also serves as the initiation site). [3] <u>upstream *eis*- *eis*</u>: the intergenic region *eis*-Rv2417c (2715333 - 2715383) for SNVs and indels, and the coding sequence for *eis* (2714124 - 2715332) for nonsense SNVs, frameshift indels, missense SNVs that abolish the start codon, and synonymous SNVs that abolish the START codon. [4] <u>upstream *whiB7*- *whiB7*</u> the intergenic region *whiB7*-*uvrD2* (3568680 - 3569082) for SNVs and indels, and the coding sequence for *whiB7* (3568401 - 3568679) for nonsense SNVs, missense SNVs that abolish the start codon, frameshift indels, and synonymous SNVs that abolish the start codon.

### Antibiotic Resistance Mutations in *rrs* and *atpE*

Resistance to aminoglycosides can occur as a result of mutations in the 1,400bp region of the 16S rRNA (*rrs*), where *rrs* A1401G, C1402T, and G1484T mutations have all been implicated in aminoglycoside resistance (Kambli et al., 2016; Reeves et al., 2013). To ensure that isolates were not aminoglycoside resistant directly from harboring one of these *rrs* mutations, we genotyped (with ≥75% allele frequency) the 1401, 1402, and 1484 nucleotide coordinates in *rrs* for the set of 12 added isolates with *eis* C-14T promoter resistance mutations and 17 other isolates (from our original set of 31428 isolates) with coinciding *eis* C-14T promoter resistance mutation and *eis* LoF mutations (**Fig. 3, Table 3, Supplementary Table 3**). None of these 29 isolates harbored any of the *rrs* A1401G, C1402T, or G1484T aminoglycoside resistance mutations (**Table 3**). Similarly, single nucleotide variants in the gene *atpE*, which encodes the BDQ target, have been associated with high-level BDQ resistance (Kadura et al., 2020). We interrogated the genotypes for 29 SNP sites in *atpE* (SNPs that were present within our pool of 31,428 isolates) in the 84 isolates that harbored both a frameshift in *mmpR* and frameshift in *mmpL5* (**Fig. 1, Table 2**) and found that none of the isolates carried a mutant allele at any of these SNP sites.

### Phylogeny Construction and assessment of convergent evolution

To generate the trees, we first merged the VCF files of the isolates in the sample (188 lineage 4.11 isolates & 444 lineage 2.2.1.1.1.3.i3 isolates) with bcftools (Li et al., 2009). We then removed repetitive, antibiotic resistance and low coverage regions (Freschi et al., 2020). We generated a multi-sequence FASTA alignment from the merged VCF file with vcf2phylip (version 1.5, https://doi.org/10.5281/zenodo.1257057). We constructed the phylogenetic tree with IQ-TREE (Nguyen et al., 2015). We used the *mset* option to restrict model selection to GTR models, implemented the automatic model selection with ModelFinder Plus (Kalyaanamoorthy et al., 2017) and computed the SH-aLRT test and bootstrap values with UFBoot (Minh et al., 2013) with 1000 bootstrap replicates.

To quantify the number of independent mutational events (SNPs & indels) in the original sample of 31,428, we grouped isolates into eight groups based off of genetic similarity, five groups corresponding to global lineages 1, 2, 3, 5, 6 and three groups for global lineage 4. We constructed eight phylogenies from these groups, then used the genotypes in conjunction with the phylogenies to assess the number of independent arisals for each mutation observed. We used an ancestral reconstruction approach to quantify the number of times each SNV arose independently in the phylogenies using SNPPar (Edwards et al., 2020). This yielded a *homoplasy score* or an estimate for the number of independent arisals for each SNV (**Supplementary Table 1**). To quantify the number of independent arisals for each indel, we developed a simple method to count the number of times each indel allele “breaks” the phylogenies. If a given mutant allele is observed in two separate parts of a phylogeny, then we can assume that this allele arose twice in pool of isolates used to construct the tree. We calculated a *homoplasy score* by counting these topology disruptions for both SNVs & indels. The results for the SNVs were congruent with the *homoplasy scores* computed from the ancestral reconstructions, validating this approach for computing *homoplasy scores* for indels.

### MRCA Dating Approximation

To date the arisal of a specific mutation within a group of isolates on a phylogeny, we looked for groups of isolates on the trees that carried the mutant allele of interest. We grouped isolates according to the following principles: (1) a group of isolates had to be a sub-tree of 2 or more monophyletic mutants, and (2) we identified the MRCA of all mutants in that sub-tree assuming that reversion of mutations is impossible. For a given group, we checked that the MRCA of the isolates had an SH-aLRT of ≥80% and an ultrafast bootstrap support of ≥95%. If these conditions were satisfied, indicating high confidence in the branch, we then calculated the median branch length (SNPs/site) between the MRCA and the tips. We multiplied the median branch length (SNPs/site) by the number of sites in the SNP concatenate used to construct the tree to get the median branch length in SNPs/genome. Molecular clock estimates for MTBC range from 0.3-0.6 SNPs/genome/year, we divided the branch lengths in SNPs/genome by 0.3 SNPs/genome/year and 0.6 SNPs/genome/year to get upper and lower bound estimates for the MRCA age.

### Data Analysis and Variant Annotation

Data analysis was performed using custom scripts run in Python and interfaced with iPython (Pérez and Granger, 2007). Statistical tests were run with Statsmodels (Seabold and Perktold, 2010) and Figures were plotted using Matplotlib (Hunter, 2007). Numpy (Van Der Walt et al., 2011), Biopython (Cock et al., 2009) and Pandas (McKinney and others, 2010) were all used extensively in data cleaning and manipulation. Functional annotation of SNPs was done in Biopython using the H37Rv reference genome and the corresponding genome annotation. For every SNP variant called, we used the H37Rv reference position provided by the Pilon (Walker et al., 2014) generated VCF file to determine the nucleotide and codon positions if the SNP was located within a coding sequence in H37Rv. We extracted any overlapping CDS region and annotated SNPs accordingly, each overlapping CDS regions was then translated into its corresponding peptide sequence with both the reference and alternate allele. SNPs in which the peptide sequences did not differ between alleles were labeled synonymous, SNPs in which the peptide sequences did differ were labeled non-synonymous and if there were no overlapping CDS regions for that reference position, then the SNP was labeled intergenic. Functional annotation of indels was also done in Biopython using the H37Rv reference genome and the corresponding genome annotation. For every indel variant called, we used the H37Rv reference position provided by the Pilon generated VCF file to determine the nucleotide and codon positions if the indel was located within a coding sequence in H37Rv. An indel variant was classified as in-frame if the length of the indel allele was divisible by three, otherwise it was classified as a frameshift.

## SUPPLEMENTARY TABLE DESCRIPTIONS

**Supplementary Table 1. Mutations detected in a global sample of MTBC clinical isolates**. A full list of mutations that occur within our sample of 31,440 clinical isolates within the *mmpL5, mmpS5, mmpR, ahpC, eis, whiB7* coding sequences and *oxyR*-*ahpC, eis*-Rv2417c, *whiB7*-*uvrD2* intergenic regions.

**Supplementary Table 2. Mixed indels in the *mmpR*-*mmpL5*-*mmpS5* chromosomal region**. A list of frameshift indels that were detected at an intermediate allele frequencies between 10% and 75% in *mmpR, mmpS5*, or *mmpL5* within our sample of 31,428 isolates (excludes the set of 12 added isolates, see **Methods**).

**Supplementary Table 3. Co-occurrence of regulator resistance mutations and regulon LoF mutations**. A more detailed version of **Table 2**.

**Supplementary Table 4. Binary resistance phenotypes for MTBC sub-lineage 4.11 isolates**. A table of binary resistance phenotype (STR, INH, RIF, EMB, PZA, AMK & KAN) data for a subset isolates that belong to sub-lineage 4.11 (**Fig. 2**), curated from multiple studies (Groschel et al., 2021).

**Supplementary Table 5. Count of isolates with *eis* promoter mutations and no coinciding *rrs* AG resistance mutations**. The count of isolates with *eis* promoter mutations (G-10A, C-12T, C- 14T, G-37T) that coincide with any AG resistance mutations in *rrs* (A1401G, C1402T, G1484T).

**Supplementary Table 6. KAN and AMK resistance details for strains with MICs and strains with double *eis* promoter SNP & *eis* LoF mutations**. A more detailed version of **Table 3** with binary resistance phenotype (STR, INH, RIF, EMB, PZA, AMK & KAN) data for a subset of isolates (Groschel et al., 2021).

## ACKNOWLEDGEMENTS

We thank Koné Kaniga and Nacer Lounis for sharing information about the C208 trial conducted by Janssen and Thomas Schön for helpful discussions regarding aminoglycoside resistance. We thank the members of the Farhat lab for helpful discussions and comments on the research project and manuscript. R.V.J. was supported by the National Science Foundation Graduate Research Fellowship under Grant No. DGE1745303. C.U.K. received an observership from the European Society of Clinical Microbiology and Infectious Diseases. M.R.F. was supported by NIH NIAID R01 AI55765. Portions of this research were conducted on the O2 High Performance Compute Cluster, supported by the Research Computing Group, at Harvard Medical School.

## COMPETING INTERESTS

C.U.K.’s work for Becton Dickinson involves a collaboration with Janssen and Thermo Fisher Scientific. C.U.K. is a consultant for Becton Dickinson, the Foundation for Innovative New Diagnostics, the Stop TB Partnership, and the TB Alliance. C.U.K. worked as a consultant for QuantuMDx, the WHO Global TB Programme, and WHO Regional Office for Europe. C.U.K. gave a paid educational talk for Oxford Immunotec. Hain Lifescience covered C.U.K.’s travel and accommodation to present at a meeting. C.U.K. is an unpaid advisor to BioVersys and GenoScreen.

## DATA AND MATERIALS AVAILABILITY

Mtb sequencing data was collected from NCBI and is publicly available. WGS data for the set of added 12 clinical *eis* C-14T mutants (**Materials and Methods**) will be uploaded to a public sequence repository upon acceptance of this manuscript for publication. All packages and software used in this study have been noted in the **Materials and Methods**. Custom scripts written in python version 2.7.15 were used to conduct all analyses and interfaced via Jupyter Notebooks. All scripts and notebooks will be uploaded to a GitHub repository upon acceptance of this manuscript for publication.

